# Multidimensional plasticity of gene expression underlying higher macrolide tolerance in saline and warm environments

**DOI:** 10.64898/2026.01.07.698155

**Authors:** Marie Rescan, Marc Dachs Rojo, Carles Borrego

## Abstract

Organisms are routinely exposed to multiple environmental stresses, increasingly intensified by human activities. While adaptation occurs over the scale of at least a few generations, phenotypic plasticity enables rapid adjustment to environmental changes. Gene expression is a central plastic trait that mediates phenotypic change, yet how synergistic or antagonistic fitness effects arise from interactions among transcriptional responses remains poorly understood. Here, we introduce a general framework to classify interactions at the gene-expression level, and discuss their evolutionary consequences in the pressing context of antibiotic resistance. We analyzed the transcriptional response of *Escherichia coli* to azithromycin (AZI) across two salinity and temperature conditions. *De novo* and antagonistic interactions were prevalent, with evidence of cross-regulation between salt and AZI. High salinity increased tolerance by two orders of magnitude and, similarly to AZI, promoted a metabolic shift from carbon to nitrogen, potentially facilitating the clearance of macrolide-induced misfolded proteins. Reduced temperature, which cancelled the salinity protective effect, enhanced carbon metabolism and counteracted this shift. Salinity additionally restored stress-response pathways, largely repressed by AZI. Third-order interactions attenuated the contribution of salinity relative to AZI, but the number of affected genes declined exponentially with interaction order, suggesting that higher-order interactions at the gene-expression level should play a minor role in the responses to multiple stressors. By modulating transcriptional responses to AZI, simple environmental parameters ultimately reshaped the adaptive landscape of antibiotic resistance, altering the spectrum of resistance mutations likely be fixed.

## Introduction

All organisms evolve within complex environments that undergo multivariate changes, exacerbated by human activities and potentially threatening populations persistence. Beyond their individual effects, environmental variables can interact non-additively [1–3], reducing our ability to predict demographic responses under multiple simultaneous stressors. Understanding population response to a changing complex environment requires to get insights into the mechanisms underlying multidimensional environmental tolerance.

Many genes are plastic, exhibiting variable expression levels in response to environmental cues [4, 5] and allowing for phenotypic adjustment, yet the broader role of the interactions at the level of gene expression remains poorly understood. Most studies considering the effect of multiple stressors on gene expression limited their analyses to each stressor’s individual effects [6–12]. Very little is known on the distribution of interactions at the gene expression level, especially whether these interactions generally amplify or buffer the individual effects of stressors, or can even restore expression levels affected by one single or two stressors [13, 14]. Few studies have concluded that combined stressors act like a totally novel stress, with interactions emerging *de novo* in genes that were not responsive to individual stressors [15, 16]. But none explored if higher-order interactions also occur at the transcriptomic level, nor investigated the role of the regulation network on the response to multiple stressors. Can synergy and antagonism at the fitness level result from changes in expression of a regulator in response to another stress?

We addressed these questions in the pressing context of environmental antibiotic resistance. Antibiotics are now ubiquitous at sub-lethal concentrations in most, if not all, surface waters [17, 18], where they co-occur with a range of fluctuating biotic and abiotic factors. Interactions between antibiotics and environmental variables modulate bacterial tolerance to antibiotics [19, 20], thereby altering the relative fitness of strains carrying antibiotic resistance genes (ARGs) and the evolutionary dynamics of environmental resistance. In bacteria, antibiotics inducing a heat-shock-like response were less deleterious at higher temperatures, and vice versa [21, 22], suggesting that two stressors eliciting similar gene expression responses tend to interact antagonistically. Conversely, stressors that induce opposing expression responses seem more likely to have synergetic effect on fitness, resulting in combined effects that are more than the sum of individual ones.

Azithromycin (hereafter AZI) is a macrolide frequently measured in wastewater [23], that blocks translocation during protein synthesis and, at high concentrations, leads to the toxic accumulation of misfolded proteins [24, 25]. Pairwise and third-order interactions among AZI, temperature, and salinity could alter by up to 100-fold the antibiotic concentration required to halve bacterial growth rate or yield [26], which was not attributable to differential degradation of AZI. While temperature and salinity may influence AZI activity directly by altering its affinity for the translation machinery, these environmental variables more likely modulate gene expression patterns affecting cell wall and membrane permeability to AZI, clearance of misfolded proteins, or the level of toxic stress resulting from their accumulation.

We quantified the distribution of pairwise and third-order interactions between these environmental variables, namely: salinity, temperature, and azithromycin (AZI), on the transcriptomic response of AZI-sensitive *Escherichia coli* populations. We provided a new classification framework for the interactions at the gene-expression level, linked them to the higher tolerance to AZI in saline and warm environment and discussed the consequences of these interactions on the evolutionary dynamics of resistance.

## Results

### Saline and warm conditions protected *E. coli* growth against azithromycin

Populations of *E. coli* were exposed to all cross-combinations of two salinities (0.085M and 0.585M), two temperatures (30°C and 25°C) and two concentrations (absence or 1µg mL^−1^) of AZI, and population sizes were monitored using flow cytometry (Figure 1A). We confirmed that high salinity reduced *E. coli* growth in the absence of AZI but offered a protection against this antibiotic, while lower temperatures decreased both growth rate and AZI tolerance [26]. Population size after 24h was divided by 4.9 by AZI at 30°C (p < 10^−6^). Cold temperature slightly intensified the effect of AZI (p = 5.52×10^−2^) but salinity nearly restored population size (multiplying final population size by 3.1 compared to AZI alone, p = 4.23×10^−4^). This protection weakened in lower temperature, as revealed by a negative third order interaction (*p* = 2.39×10^−3^).

**Figure 1.**
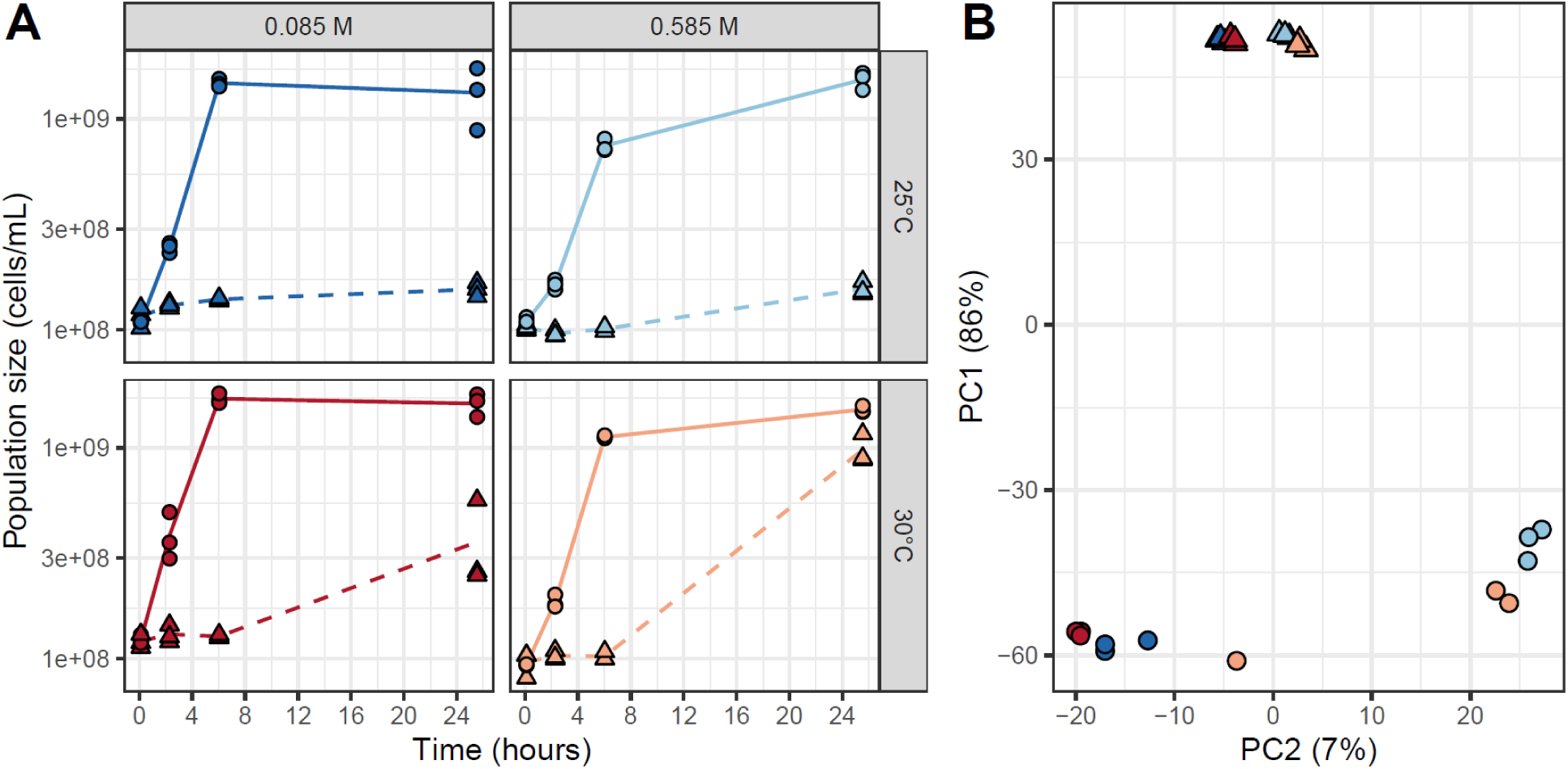
Population dynamics and transcriptomics response to combined stressors. A. *E. coli* growth after exposure to two temperatures (rows, blue *vs*. red), two salinities (columns, light *vs.* dark colors) and exposed to none (plain lines and dots) or 1 µg mL^−1^ (dashed lines and triangles) of AZI. B. Principal component analysis (PCA) of gene expression level for *E. coli* 6h after exposure to the combined treatments.

To link the interactions at the fitness level to interactions at the gene expression level, we analyzed the transcriptomes of the 24 populations, 6h after exposure. The changes in gene expression correlated well with the level of each stress. AZI, near the MIC at 25°C in BHI (Figure 1A), explained the major part of the variance in gene expression patterns (∼86%, PCA axis 1, Figure 1B). *E. coli* could tolerate up to twice the salinity used here [26], and in the absence of antibiotic, populations were clearly well-split by salinity along the second PCA axis (up to 7% of the variance). A 5°C decrease did not represent a significant stress considering that 30 °C was already suboptimal, and the temperature treatments, although well clustered, explained only a marginal part of the variance in gene expression. Although interactions with salt and temperature importantly reduced the fitness effect of AZI, the PCA ordination did not evidenced such interactions.

### Gene expression overlaps between AZI, temperature and salinity

#### Common down-regulation are more frequent than common up-regulation

We analyzed differentially expressed genes (|log2FoldChange| > 2 and adjusted *p*-value < 10⁻^6^) in populations exposed to each treatment individually. AZI induced significant expression changes in 1336 genes (651 upregulated / 675 downregulated), salinity in 111 genes (30 up / 81 down) and temperature in 35 genes (14 up / 21 down). All stressors produced more down- than up-regulation. AZI, temperature and salinity collectively induced the downregulation of the same 10 genes, with additional 37 genes commonly downregulated by AZI and salt. By contrast, no genes were commonly upregulated by the three variables (Figure 2A).

**Figure 2.**
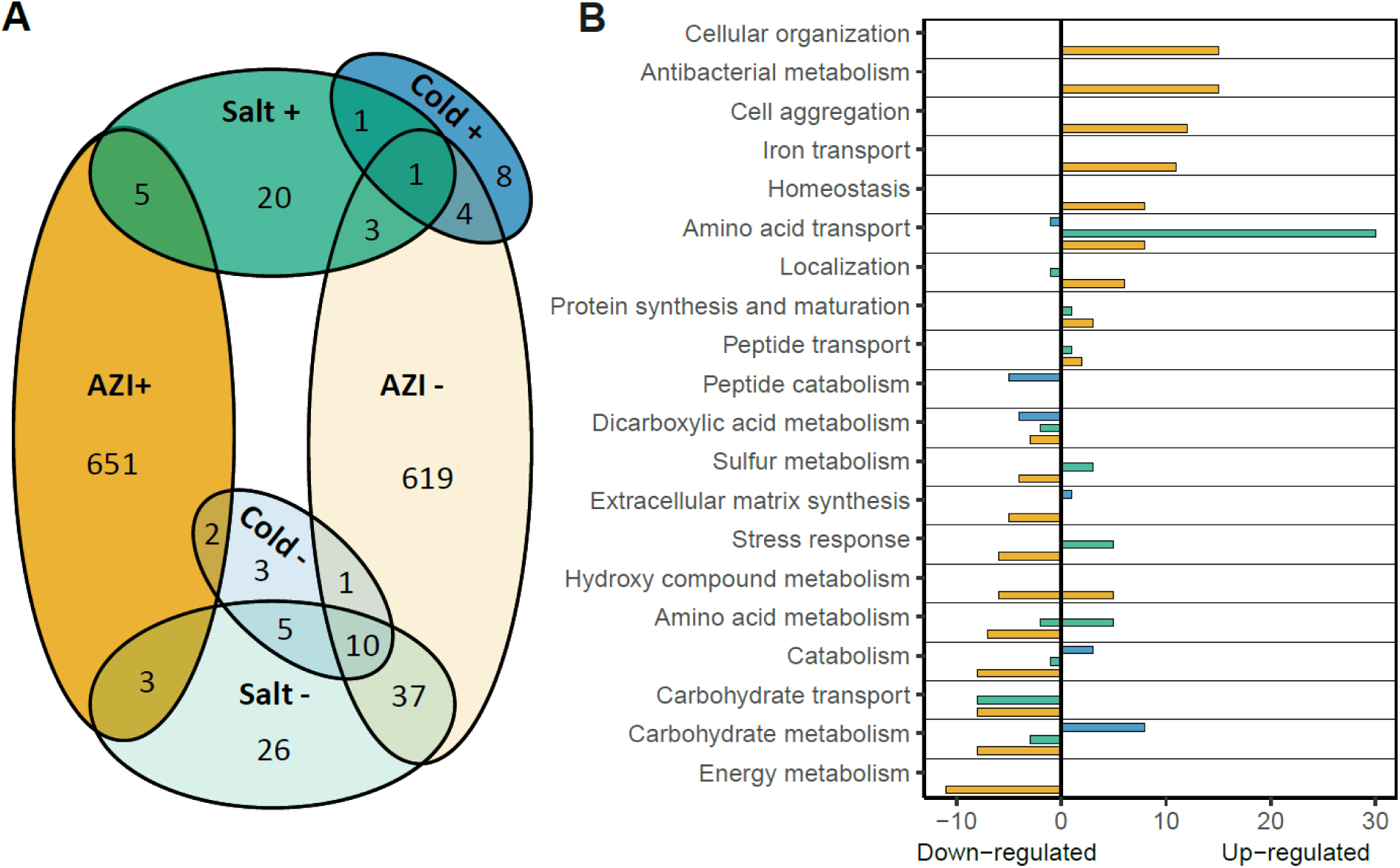
Overlaps between differentially expressed genes under experimental treatments. A. Venn diagram displaying the number of gene transcripts under-expressed (light colors) and over-expressed (darker color) 6 h after exposure to either AZI (orange), high salinity (green) or temperature (blue). B. Number of Gene Ontology (GO) terms enriched for 21 functional groups. Colors indicate the environmental variable triggering differential gene expression (orange: AZI, green: salt, blue: temperature).

#### Overlaps involved stress response and balance between nitrogen/carbon metabolism

Differentially expressed genes were functionally annotated (Figure 2B). AZI upregulated pathways related to cellular organization, antibacterial metabolism, and iron transport, supporting the idea that antibiotic resistance can be linked to metal-tolerance mechanisms [27]. It also enhanced cell aggregation, a strategy that reduces local antibiotic concentration, and upregulated homeostasis. AZI shifted the carbon–nitrogen metabolic balance by increasing amino acid transport and protein-synthesis pathways while downregulating carbohydrate transport and metabolism, energy metabolism, and amino acid catabolism. Unexpectedly, it repressed stress-response and sulfur-metabolism pathways, protective against oxidative stress [28].

Salinity effects were similar to AZI on several metabolic pathways (Figure 2B). They both enhanced amino acid transport and downregulated carbohydrate transport and metabolism. But contrary to AZI, salinity upregulated osmoprotection and sulfur metabolism, which can buffer AZI-induced ROS [29–31]. Lower temperature, that amplified the effects of AZI (Figure 1A), largely opposed AZI’s transcriptional effects, up-regulating carbohydrate metabolism and downregulating peptide transport and metabolism.

### Antagonistic and *de novo* interactions acting on metabolism and stress response further explain interactions at the fitness level

In addition to the single effect of each stressor, our set-up allowed to quantify pair-wise and third order interactions effects on gene expression. For each pair of stressors, additivity was defined by a non-significant interaction at the logarithmic scale (thick black dotted lines, Figure 3A). Genes that responded neither to the individual variables nor to their interactions were not counted. About 10% of the plastic genes displayed a non-additive response to the pairs of environmental stressors, with 282 interactions found among at least one pair of stressors, which was much higher than the 64 overlaps found between at least two single variables (Figure 2A).

**Figure 3.**
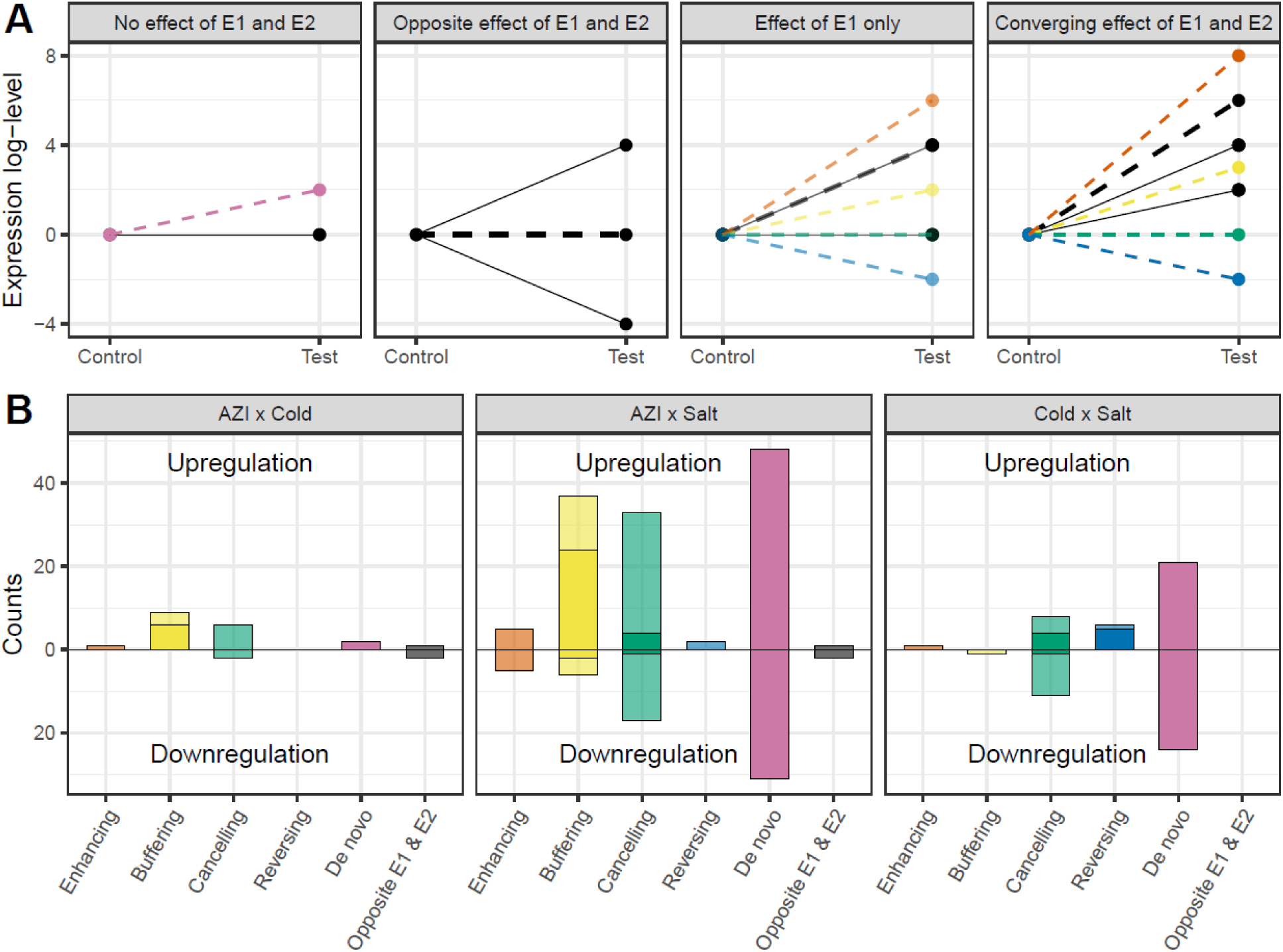
Distribution of interaction among pairs of stressors effect at the gene expression level. A. Reactions norms explaining the effect of the interaction between two environmental variables (E1 and E2, black) on gene expression. Compared to additivity (black dashed line), interactions can generate *de novo* effects (pink), enhance (red), buffer (yellow), cancel (green) or reverse (blue) the effect of E1 and E2. B. Interaction types were quantified for all combinations of pairs of stressors and split according to their regulatory effect (up or down-regulation). Bar transparency shows interactions occurring when only one (transparent) or both (plain) environmental variables influenced gene expression.

#### De novo and antagonistic were the most frequently interactions found among pairs of stressors

Interactions between two environmental stressors can either align with or counteract the sum of their individual effects on gene expression. By considering both the direction (null, up- or down-regulation) of the isolated stress effects and the sign of the interaction, interactions were classified as either “*de novo*”, “antagonistic” or “enhancing”.

*De novo* interactions, arising while both individual stressors had no significant effect, were surprisingly frequent (Figure 3, pink), and were even predominant for AZI × Salt and Salt × Temp. stressor pairs. Only few enhancing interactions (*i.e.*, that increased the additive effect of two stressors) were detected and they occurred only when only one of the stressors influenced gene expression (Figure 3, red).

Many interactions were antagonistic, counteracting the effect of either one or both environmental stressors. To dig into the effects of antagonistic interactions, we considered the sign of the individual variable effects, the sign of the interaction, and the overall effect of both interacting stressors. An antagonistic interaction that did not reverse the overall direction of the combined effect relative to the sum of the individual effects was classified as "buffering" (Figure 3A, yellow).

This applies, for instance, to genes upregulated by AZI and/or salinity, where a downregulating interaction occurs but where the overall effect remains significantly positive. These buffering interactions were identified in 43 genes for the AZI × Salt interaction, 9 genes for AZI × Temp., and only 1 gene for Temp. × Salt. Positive interactions buffering downregulation by individual variables were slightly more frequent than negative interactions (Figure 3, yellow), consistent with the fact that single stress triggered more down- than up-regulation. Interactions that cancelled out the individual effects of both stressors were classified as "cancelling" interactions (Figure 3, green). Cancelling effects were slightly more frequent than buffering interactions, affecting 50 genes for AZI × Salt, 8 for AZI × Temp., and 19 for Temp. × Salt, and again, were more frequently counteracting a downregulation of the genes, generally triggered by AZI alone. In rare cases, the interaction even led to a "reversing" effect, where the combination of both stressors induces an overall regulation in the opposite direction compared to their individual effects (Figure 3, blue). This occurred only in genes that were initially downregulated (6 genes for Temp. × Salt; 2 for AZI × Salt).

Finally, only 6 interactions occurred between two environmental stressors that had opposite effects (grey bars in Figure 3B), suggesting that antagonistic stressors were much less prone to interact.

#### Pairwise interaction with salt or temperature tend to buffer individual effects of AZI

To investigate the importance of interactions between pairs of environmental stressors and their potential impact on antibiotic tolerance, each gene was assigned to at least one of the functional groups previously defined based on its associated GO terms (Figure 4). *De novo* interactions occurred across almost all functional groups that involved in the response to single stress, as do

**Figure 4.**
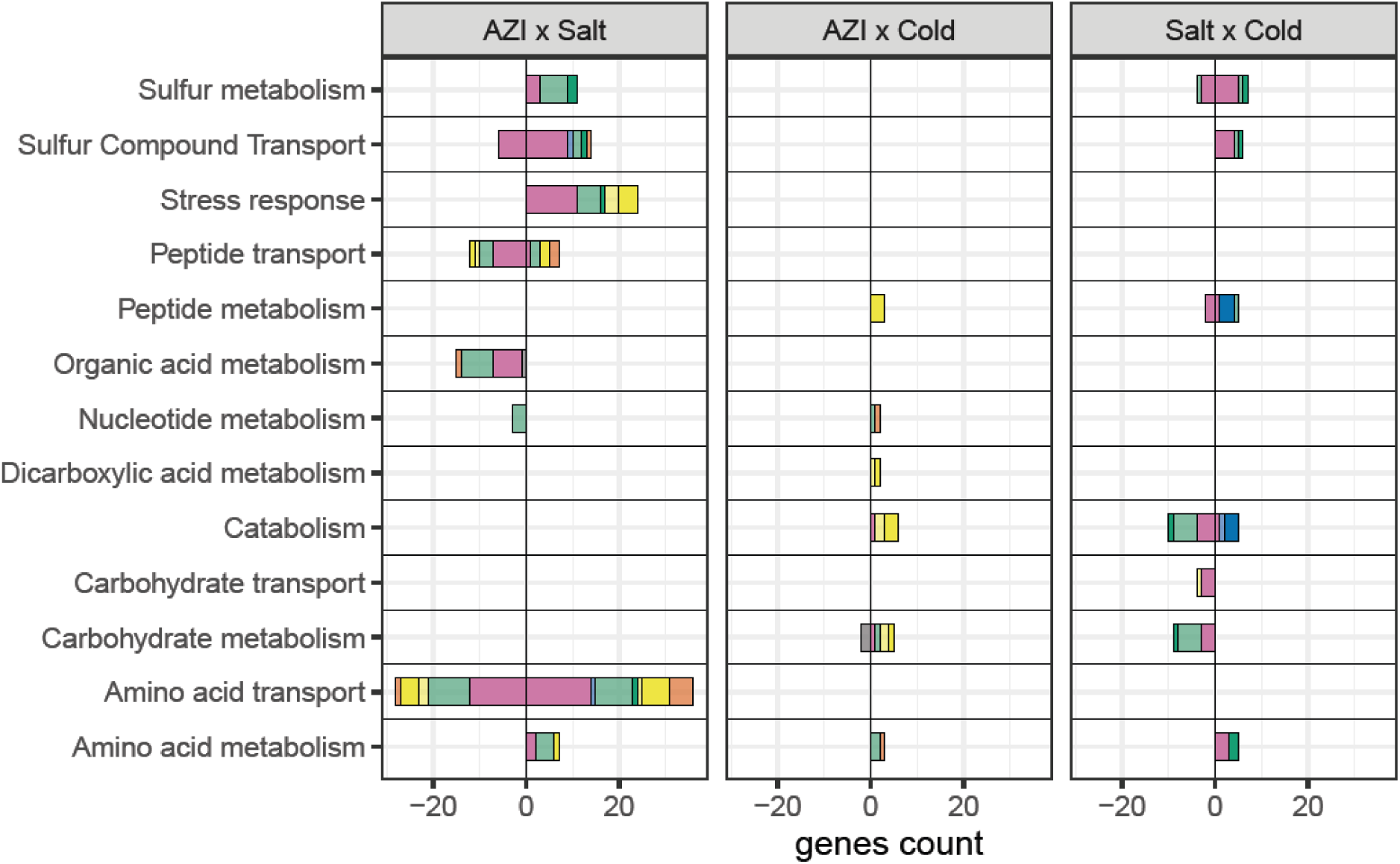
Number of genes exhibiting an interaction effect for each functional group. Colors indicate the interaction type (*de novo*: magenta, enhancing: red, buffering: yellow, cancelling: green, reversing: blue) and transparency shows interactions occurring when only one (transparent) or both (plain) environmental variables alone had an effect on gene expression.

buffering and cancelling interactions (Figure 2 and 4). Enhancing and reversing interactions were primarily associated with amino acid and peptide transport and/or metabolism.

Interactions between AZI and salt upregulated pathways related to stress response, sulfur compound transport, and metabolism, that likely play a role in AZI tolerance, and upregulated amino acid metabolism. These groups were up-regulated by salinity and down-regulated by AZI (Figure 2), such that at the functional level, AZI × Salt interactions reinforced salt protective effect on expression. Very few interactions occurred between temperature and AZI, and they were mostly positive. At the functional level, these interactions further amplified the antagonism between AZI and low temperatures, activating expression in genes within functional groups such as carbohydrate metabolism or catabolism, that were downregulated by AZI (Figure 2 and 4). More interactions were observed between temperature and salinity than between temperature and AZI, despite the fact that AZI regulates about 10 times more genes than salinity. They enhanced expression of genes related to sulfur compound transport and reduced those involved in carbohydrate metabolism and transport.

*Few third-order interactions explain weaker salinity protection in cold*.

Forty-one genes exhibited third-order interactions among AZI, salinity and temperature. We observed a near-perfect exponential decline in the average number of genes differentially expressed with increasing interaction order (R² > 0.98, Figure 5). Although the number of stress conditions was limited, such trend suggests that higher order interactions (>3) should be rare and play only a marginal role on the response to multiple stresses.

**Figure 5.**
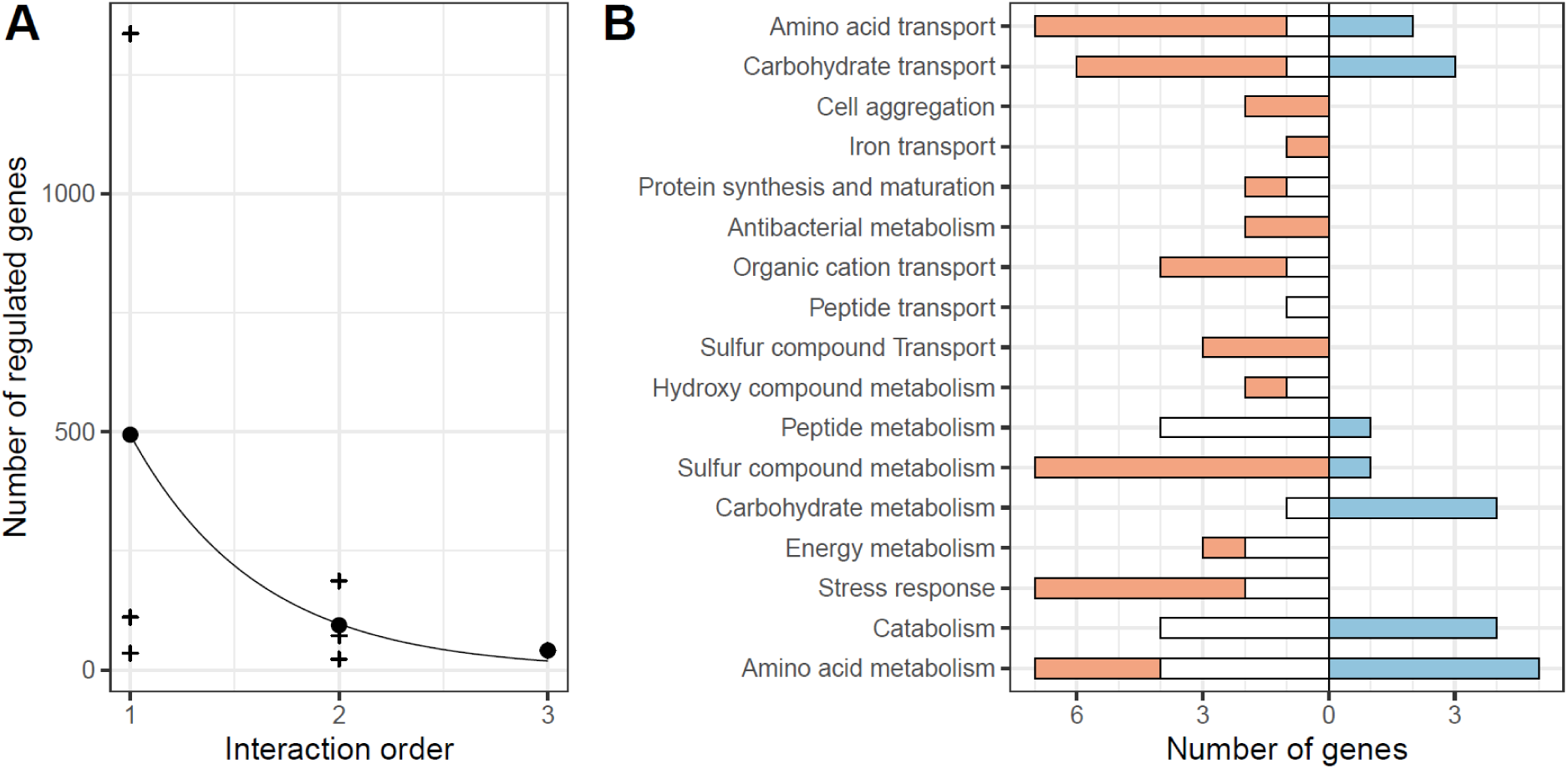
**Third order interactions**. A. Number of genes affected by the order of the interaction. For order 1, the effect of each environment alone was considered. Crosses indicate the counts for each interaction order and each environment while dots show the mean count for each interaction order. The line corresponds to a Poisson regression. B. Number of genes up- (right) or down- (left) regulated by the third order interaction in each functional semantic group. Colors indicate the overall log2 fold change induced before accounting for the third order interaction (negative < –2: blue; positive > 2: orange; null, within [–2,2]: white).

To classify the nature of the three-way interactions as enhancing, *de novo*, or antagonistic, we compared the predicted additive effect of individual stresses and pairwise interactions to the direction of the third-order interaction. Significant interactions were classified as *de novo* when the predicted sum for the three variables and their pairwise interactions was within [–2, +2], but as no statistical significance was inferred for the prediction without third-order interaction, the number of *de novo* third-order interactions was likely underestimated. All positive third-order interactions were antagonistic, while negative third-order interactions were either *de novo* or antagonistic. No enhancing third-order interactions were observed (Figure 5).

In many functional groups directly linked to antibiotic tolerance – antibacterial metabolism, iron transport, cell aggregation, sulfur compound transport and stress response – third-order interactions led to the downregulation of genes that were either upregulated or not differentially expressed in response to individual stressors and their pairwise interactions. Such downregulation driven by third-order interactions particularly counteracted the effect of salinity, explaining its reduced protective effect in colder environments.

### Regulation network response to single and multiple stressors

We investigated whether the interactions observed between different environmental stressors were shaped by the gene regulatory network, and specifically, if differential expression in response to AZI could be influenced by the response of regulators to temperature or salinity. Regulator and target genes responses to each single stressor were summarized in Table 1 and multinomial logistic regressions were performed to assess the influence of regulator gene responses on the responses of their target, in reaction to the same, or a different stressor.

**Table 1.** Stressors effect on pairs of regulator–regulated genes. . For each stress, pairs of regulator-regulated genes were counted depending on the differential expression of the regulator (columns), the regulated (rows), and on the sign of the regulation (R for repressor, E for enhancer). Regulators responding to salt and tenperature were only over-expressed.

|  |  |  | Regulator response to: |  |  |  |  |  |  |  |  |  |  |  |  |  |
| --- | --- | --- | --- | --- | --- | --- | --- | --- | --- | --- | --- | --- | --- | --- | --- | --- |
|  |  |  | AZI |  |  |  |  |  | Salt |  |  |  | Cold |  |  |  |
|  |  |  | down |  | 0 |  | up |  | 0 |  | up |  | 0 |  | up |  |
|  |  |  | R | E | R | E | R | E | R | E | R | E | R | E | R | E |
| Target response to: | AZI | down | 106 | 254 | 451 | 905 | 37 | 23 | 594 | 1179 | 0 | 3 | 594 | 1180 | 0 | 2 |
|  |  | 0 | 328 | 600 | 1207 | 2426 | 71 | 96 | 1601 | 3120 | 5 | 2 | 1606 | 3122 | 0 | 0 |
|  |  | up | 87 | 88 | 274 | 493 | 37 | 24 | 394 | 605 | 4 | 0 | 398 | 605 | 0 | 0 |
|  | Salt | down | 7 | 63 | 52 | 117 | 7 | 1 | 66 | 178 | 0 | 3 | 66 | 181 | 0 | 0 |
|  |  | 0 | 496 | 864 | 1843 | 3631 | 137 | 141 | 2470 | 4634 | 6 | 2 | 2476 | 4634 | 0 | 2 |
|  |  | up | 17 | 8 | 32 | 63 | 0 | 1 | 46 | 72 | 3 | 0 | 49 | 72 | 0 | 0 |
|  | Cold | down | 4 | 15 | 13 | 25 | 0 | 0 | 17 | 40 | 0 | 0 | 17 | 40 | 0 | 0 |
|  |  | 0 | 506 | 911 | 1897 | 3754 | 143 | 143 | 2537 | 4803 | 9 | 5 | 2546 | 4806 | 0 | 2 |
|  |  | up | 10 | 7 | 16 | 23 | 0 | 0 | 26 | 30 | 0 | 0 | 26 | 30 | 0 | 0 |

#### Single stress regulation

Reduced temperature activated one single regulator (*metR*), but none of its two targets displayed differential expression in response to temperature. The response to salt also involved the upregulation of a single regulator, *stpA*. Genes were more likely downregulated in response to salt when they were activated by *stpA* (*p* = 3.74×10⁻⁵) and conversely, genes upregulated in response to salt were significantly overrepresented among those repressed by *stpA* (*p* = 1.67×10⁻⁶), highlighting the stabilizing role of this regulator [32]. The response to AZI involved 672 regulator–target pairs, involving 45 regulators and 177 target genes both showing significant differential expression (Figure 6). Genes under-expression in response to AZI did not significantly depend on the response of their regulators (*p* > 0.12). Target genes upregulated in response to AZI were overrepresented among those whose repressors were also upregulated (*p* = 1.22×10⁻⁵) and underrepresented among those whose activators were downregulated (*p* = 2.58×10⁻³), suggesting again a canalization of gene expression [33]. Major regulators included the global transcriptional activator *crp*, a system response repressor *arcA*, and regulators involved in stress response (*cpxR*), acid stress response (*ydeO* and *gadE*) and flagellar transcription (*flhC* and *flhD*). The differentially expressed target genes were typically regulated by a limited number of regulators (≤ 7). Among the most highly connected targets were genes encoding proteins involved in biofilm formation (*e.g. csgF*, *csgG*), the acid resistance transcriptional activator *gadE*, and a subunit of a multidrug efflux transporter *mdtE*.

**Figure 6.**
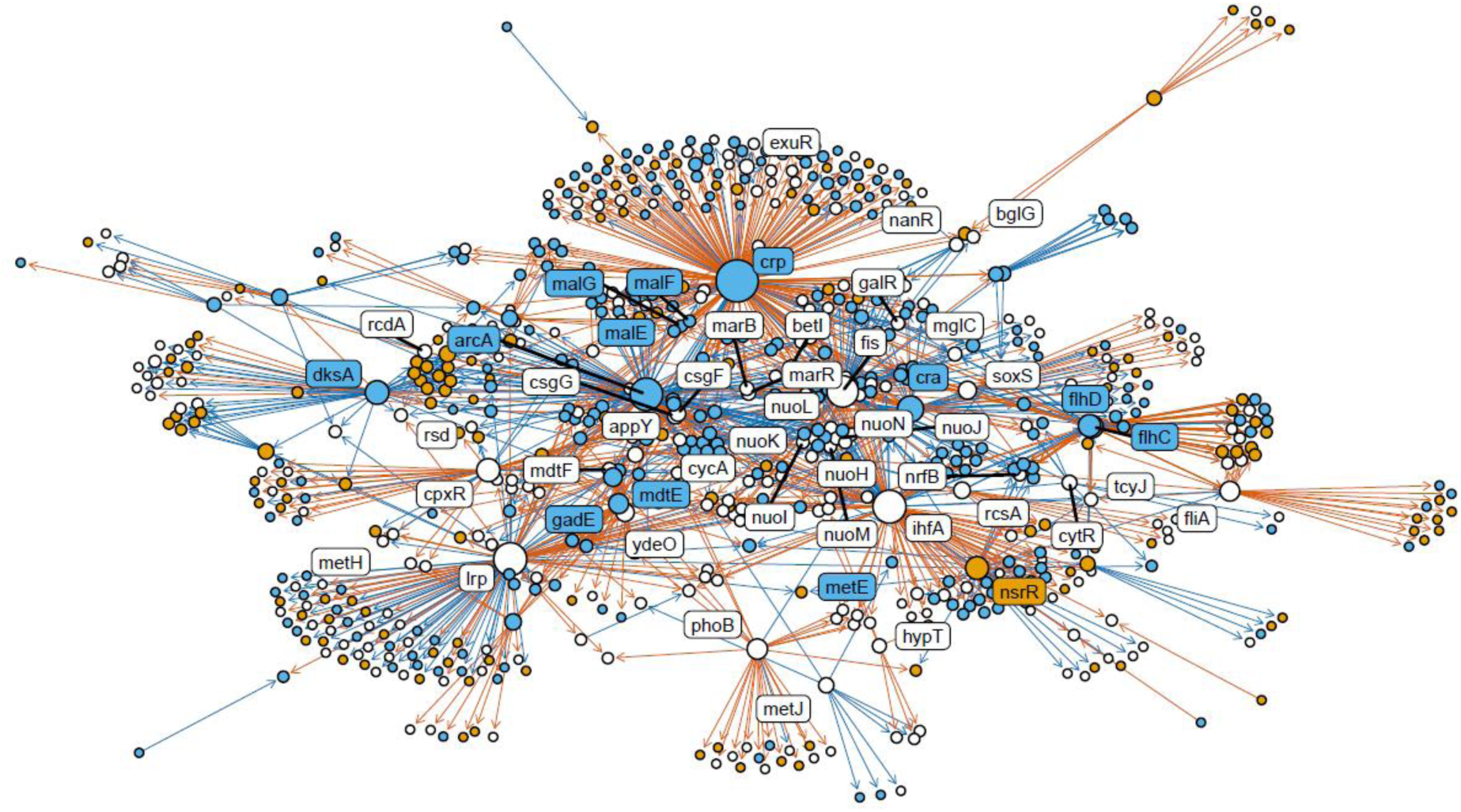
Regulation network response to AZI stress under different salinity and temperatures. The node color indicates the sign of gene expression change in response to AZI (orange: always overexpressed, blue : always underexpressed, white: depends on temperature and salinity conditions). The arrow color indicates the sign of the regulation (blue: repressed, orange: enhanced). Genes that did not significantly responded to AZI are not displayed. Labels identify genes that are highly connected (> 20 connections for constant direction of expression change – blue or orange, > 4 for varying direction – white) or that are highlighted in the main text.

#### Genes network response to AZI depended on salinity and temperature

We compared gene expression levels in response to AZI within the regulatory network under different temperature and salinity conditions. To this end, we performed independent differential expression analysis contrasting each of the four AZI treatments with the control treatment (30 °C, minimal salinity, no AZI). Within the network of genes responsive to AZI in at least one condition, 38% of the responses qualitatively changed, between up-regulated, down-regulated and NS, with temperature and salinity. This ratio was 37% for the targets and 46% for the regulators, that were therefore more prone to respond to environmental variations. The regulators which differential expression in response to AZI depended on salinity and temperature were involved in stress response (*cpxR*), flagellar biosynthesis (*fliA*), global transcriptional regulation (*lrp* and *fis*), and stress response (*phoBO* and *soxS*). The target genes displaying environment-dependent changes in expression were distributed evenly across the network (Figure 6).

Salt and lower temperature affected respectively only one regulator, not responsive to AZI (*stpA* and *metR*). Genes upregulated by AZI were significantly overrepresented among genes inhibited by *stpA* (p = 4.77×10⁻²) and underrepresented among genes activated by *stpA* (p < 1×10⁻^8^), suggesting that cross-regulation participate in the antagonism between salinity and AZI. By contrast, no significant relationship was found between genes response to AZI and their regulation by *metR* (p = 5.59×10⁻²).

## Discussion

We investigated how interactions among transcriptomic responses shape fitness-level interactions in the context of antibiotic tolerance in variable temperatures and salinities. Expression patterns induced by AZI, temperature, and salinity in isolation provide first insights into fitness-level interactions. AZI near the MIC reprogrammed almost one-third of all genes, many linked to AZI tolerance. Antibacterial metabolism and genes involved in homeostasis and cell aggregation, that reduce local AZI concentration [34], were enhanced. Iron transport was upregulated, consistent with cross-resistance involving metal–antibiotic efflux pumps [20, 35, 36]. AZI also triggered a metabolic switch from carbon to nitrogen resource, that likely contribute in eliminating and recycling the misfolded proteins accumulating in the cytoplasm due to the impaired protein synthesis caused by macrolides [24, 25].

Salinity provided a protective effect against AZ and induced similar effects on metabolism, while reduced temperatures, which exacerbated the detrimental effects of AZI, upregulated carbohydrate and downregulated peptide metabolism. Salinity additionally enhanced stress responses, especially sulfur metabolism, that were down-regulated by AZI. H2S plays a well-documented protective role against oxidative stress [29, 31], such that under high salinity, sulfur metabolism pathways can participate to antibiotic tolerance [28, 30, 37, 38].

Overall, 10% of genes exhibited pairwise interactions, with 38% of the network of genes involved in AZI response displaying a different qualitative response depending on temperature and salinity. Pairwise interactions were predominantly *de novo* or antagonistic, sometimes even abolishing single-stressor effects, while very few interactions amplified nor reverted the effects of one or both environmental variables. AZI × temperature interactions were limited, whereas AZI × salt interactions were numerous: they upregulated stress-response genes and further promoted nitrogen over carbon metabolism, reinforcing salinity’s protective effect. D*e novo* interactions were the most frequent, but occurred exclusively within functional groups already altered by single stressors. At the functional level, *de novo* interactions tended to counteract the effects of AZI, consistent with the predominance of antagonistic interactions observed at the gene level. Very few significant interactions emerged when two environmental variables exerted opposing effects on gene expression. While interactions mostly restored baseline expression levels, it seems that when additivity alone allows to stay close to this level, combined stress do not induce significant interactions.

Third-order interactions among AZI, salinity, and temperature affected a minor fraction of genes but yet tended to suppress salinity’s protective effect by downregulating antibacterial metabolism, iron transport, stress responses, and cell aggregation, while shifting energy metabolism toward carbon. This explains why salinity provided a weaker protection against the detrimental effects of AZI at lower temperature. The number of significant interactions decreased exponentially with increasing interaction order, suggesting that higher-order interactions rapidly become negligible in the context of multiple stressors. Finally, while our conclusions are broadly generalizable, the identity and direction of antibiotic × environment interactions are likely species-specific, depending on each species’ thermal and salinity niche [26].

Environmental antibiotic concentrations are far below the MIC of most microorganisms, yet they still impose a continuous selective pressure promoting the evolution of resistance [39, 40]. By modulating antibiotic tolerance, temperature and salinity can alter the evolutionary trajectory of resistance. In heavily contaminated environments with initially sensitive bacterial populations, warmer and more saline conditions may facilitate resistance emergence because enhanced tolerance enables populations to survive longer, increasing the likelihood that resistance-conferring mutations arise and spread. Conversely, in urban wastewater systems, where antibiotic levels are typically sub-MIC and ARGs are continuously introduced via human feces [41, 42], higher tolerance decreases the difference in growth rate between sensitive and resistant strains, reducing the selective advantage of resistant strains, thereby limiting their dissemination. Regulation of carbon and nitrogen metabolism emerges as a central factor in these interactions, and nutrient availability is likely an additional determinant of the dynamics of resistance.

Transcriptomic analyses revealed that multiple pathways contributing to AZI tolerance can be either activated or repressed depending on temperature and salinity. Because gene expression positively correlates with selection strength [43, 44], a mutation conferring resistance should fix more rapidly under antibiotic pressure if salt, temperature, or their interaction with AZI drive its overexpression. Conversely, mutations that confer resistance but carry fitness costs in the absence of antibiotics are more likely to persist at low frequency via genetic drift if they occur in genes under-expressed and exposed to weaker purifying selection. Thus, even common environmental parameters such as temperature and salinity can reshape the adaptive landscape of antibiotic resistance, altering not only the dynamics of resistance evolution but also the spectrum of resistance mutation that are likely to spread.

## Methods

### Strain and environmental treatments

*Escherichia coli* K12 MG1665 was cultured in Brain Heart Infusion (hereafter BHI) and exposed to all combinations of two salinities (0.085 M and 0.585 M NaCl), two temperatures (25°C and 30°C) in the presence or absence of azithromycin (1µg mL^−1^). Each treatment was replicated 3 times, making 24 independent populations.

Cultures were initiated from a frozen stock, plated on BHI agar and grown for 3 days at 30°C. One colony was then picked and inoculated into 10 mL BHI broth. After overnight growth, cultures were diluted by 1:200 into fresh BHI broth in two Falcon Tubes placed overnight at the treatment temperature (25° and 30°C). Cultures were then diluted in fresh BHI to reach an OD590 = 0.56, and 500 µL of this pre-diluted culture was transferred into 6-well microplates filled with 6.5 mL of BHI with targeted salinity and antibiotic concentrations and incubated for 3 days.

### Population density measures

Population sizes were assessed by flow cytometry, right after inoculation and after 2h, 6h and 24h. Cultures were mixed manually by carefully pipetting up and down, and 10 µL were sampled and diluted from 10^−4^ to 10^−5^ into sterile PBS, stained with the nuclear binding fluorescent dye Cystain® Green and analyzed through a Cube6^®^ flow cytometer (Sysmex, Germany). Resulting files from cytometric measurements were processed using the Floreada.io on-line tool (https://floreada.io/). Gating was performed based on side scatter (SSC) and green fluorescence (FL1) to separate *E. coli* cells from noise and debris (Supp. Figure 1). The density of events in the gate was corrected by the dilution factor to compute population density.

### RNA extraction, library preparation and Illumina sequencing

After 6h, microplates were vortexed and 1.5 mL were harvested from each of the 24 cultivation wells. Culture samples were then diluted in 3 mL RNAprotect (QIAGEN, Germany), mixed for 5s and incubated for 5 min at room temperature. The samples were centrifugated at 5000 × *g* for 20 min, the supernatant was removed, and pellets frozen at –80°C.

RNA was extracted and purified using the RNeasy (R) Protect Bacteria Mini Kit (QIAGEN, Germany) with a previous digestion with lysozyme and proteinase K. Briefly, pellets were lysed during 10 min with agitation in TE buffer containing 20 mg mL^−1^ proteinase K and 15 mg mL^−1^ lysozyme. The lysate was washed on RNeasy Mini spin columns with RLT containing 10 µL mL^-1^ β-mercaptoethanol. Ethanol was added, and the mix was vortexed after each step. Solutions resulting from extraction were washed twice on RNeasy Mini spin columns with buffer RW1 then RPE, and RNA was finally eluted in 50 µL RNase-free water.

RNA was quantified using the Qubit RNA Assay Kit on a Qubit 4 fluorometer (ThermoFisher Scientific^®^), and RNA integrity as assessed by an electrophoreses on a 3.7% bleach agarose gel [45] with SYBR Safe DNA gel Stain (ThermoFisher Scientific^®^). RNA quantity was lower than the sequencing threshold for two replicates of the environmental treatment at 30°C, [NaCl] = 0.085 M and [AZI] = 0. To overcome this limitation, we split the remaining replica population for this treatment in 3 subsamples that were further subjected to RNA extraction as explained above. The three resulting RNA extracts were hereafter treated as independent replicates despite being pseudoreplicates for this treatment conditions.

RNA extracts were sent to the National Center for Genomic Analysis (CNAG, Barcelona, Spain) for library preparation with the Ribo-Zero Plus Microbiome Depletion kit for ribosomal RNA removal, and Illumina NovaSeq 6000 sequencing of 50 bp paired-end strands in two lanes.

### Read counts

Reads from the first and second lanes were merged, and sequence quality was checked using FastQC [46] and MultiQC [47]. Trimmomatic [48] was used for Nextera adapter removal, trimming of the 3 first three bases with medium quality, and a sliding window of 5 bases was used to remove all reads with an average score quality below 25 in the window. Curated reads were then aligned and mapped to the *E. coli* K12 MG1665 reference genome (NCBI GCF_904425475.1) using HiSat2 [49] with parameters optimized for bacterial genomes (non-spliced alignment and retaining up to 200 alignments per read to handle repetitive regions). SAM files were then converted to BAM format, sorted, and indexed with SamTools [50]. Read counting was performed using FADU. Raw fastq. and processed files were deposited on the NCBI Gene Expression Omnibus (GEO) repository with accession number GSE308980.

### Differential expression analysis

The following analyses were performed on R [52] and the code deposited in ZENODO under accession number 17493233.

Differential expression was performed using the DESeq2 package [53]. Raw counts from FADU were imported and organized into a count matrix. Metadata including salinity, temperature and antibiotic concentration were provided for all samples. Eq. 1 was used to test the effect of the interactions between salinity, temperature and AZI, with each explanatory variable treated as a factor.

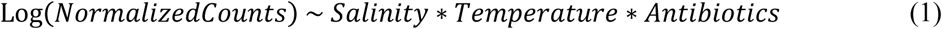

Genes were considered significantly expressed for *p*-values < 10^−6^ and log fold change below –2 or above 2.

Significant interactions were then classified depending on their effect on the expression level. Each interaction occurring between two environmental variables triggering the same effect (down-regulation or up-regulation) or no effect for one of them, was classified between antagonistic (opposite effect as the single variables in isolation) or synergetic (same effect). Antagonistic interactions were further classified as buffering, reversing or cancelling, if the global change in gene expression was conserved, cancelled or reversed compared to the additive effect of the variables in isolation. This global expression was tested using Eq. 2, where treatment correspond to a combination of antibiotic, temperature and/or salinity and was tested against the control treatment (30°C, [NaCl] = 0.085M, and [AZI] = 0).

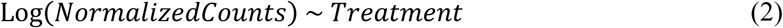

### Gene ontology and semantic analysis

All differentially expressed genes were annotated using the NCBI reference GCF_904425475.1 used for mapping. Gene ontology (GO) enrichment analysis was performed to identify biological processes significantly enriched under the different treatments and *p*-values obtained from the differentially expression analysis (Eq. 1). We annotated the enriched GO terms by performing a semantic analysis. All enriched GO terms were clustered by kmeans. Sets of clusters were defined for 10 to 30 centers. Silhouette width was higher for 24 centers, and 24 clusters were therefore kept for biological interpretation.

### Gene regulation network

*E. coli* gene regulation network was downloaded from Ecocyc [54]. Overall, 9,440 interacting pairs of regulator/target were labelled as “activating” or “inhibiting” and the regulator and target response to AZI, salt and temperature and their interactions was established from the differential expression analysis (Eq. 1). Multinomial logistic regressions were performed to assess the influence of regulator gene responses on the responses of their target genes, in reaction to the same, or a different stressor. Finally, 1238 pairs of regulator/target genes both differentially expressed under AZI in at least one temperature and salinity conditions (Eq. 2) were kept for designing the AZI regulation network and its sensitivity to the environmental conditions.

## Acknowledgment

Authors thank Arnaud Le Rouzic and Appoline Petit for their constructive suggestions.

