## Supplementary figures and images for "Multidimensional plasticity of gene expression underlying higher macrolide tolerance in saline and warm environments"

### Supp. Figure 1

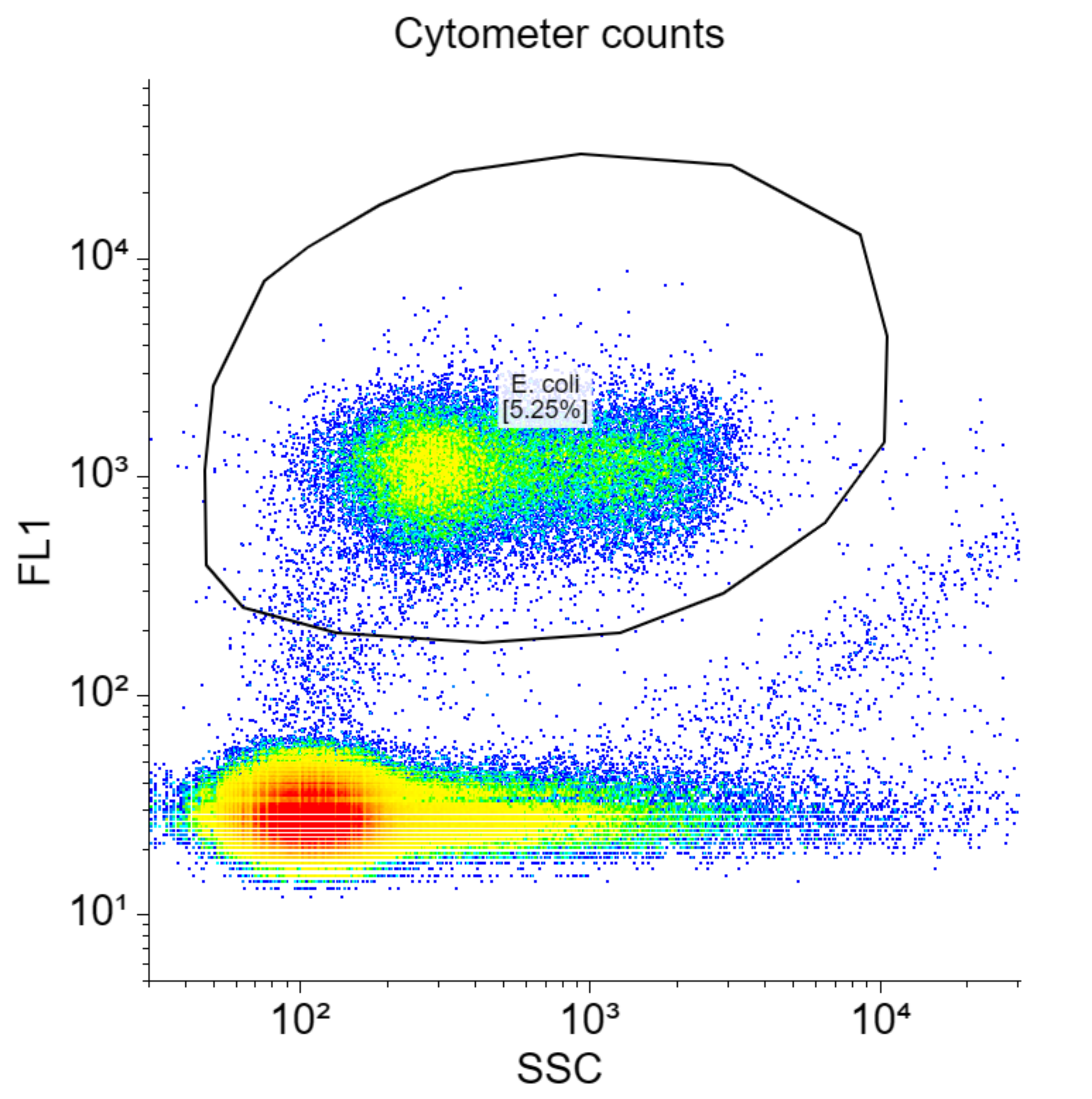
